# Detection of specific uncultured bacteriophages by fluorescence *in situ* hybridisation in pig microbiome

**DOI:** 10.1101/2022.02.18.481118

**Authors:** Line Jensen Ostenfeld, Patrick Munk, Frank M. Aarestrup, Saria Otani

## Abstract

Microbial communities have huge impacts on their ecosystems and local environments spanning from marine and soil communities to the mammalian gut. Bacteriophages (phages) are important drivers of population control and diversity in the community, but our understanding of complex microbial communities is halted by biased detection techniques. Metagenomics have provided a method of novel phage discovery independent of *in vitro* culturing techniques and have revealed a large proportion of understudied phages. Here, five large phage genomes, that were previously assembled *in silico* from pig faecal metagenomes, are detected and observed directly in their natural environment using a modified phageFISH approach, and combined with methods to decrease bias against large phages. These phages are uncultured with unknown hosts. The specific phages were detected by PCR and fluorescent *in situ* hybridisation in their original faecal samples as well as across other faecal samples. Co-localisation of bacterial signals and phage signals allowed detection of the different stages of phage life cycle. All phages displayed examples of early infection, advanced infection, burst, and free phages. To our knowledge, this is the first detection of jumbophages in faeces, which were investigated independently of culture, host identification, and size, and based solely on the genome sequence. This approach opens up opportunities for characterisation of novel *in silico* phages *in vivo* from a broad range of gut microbiomes.

## Introduction

Viruses of bacteria (phages) are an important part of microbial communities, including human and animal gut microbiomes (Brüssow & Hendrix, 2002; Shkoporov & Hill, 2019). The gut microbiome has been shown to affect the host in many aspects from nutrient uptake (Quan et al., 2019; Yang et al., 2017) to diabetes (Li, Stirling, Yang, & Zhang, 2020), obesity (Guirro et al., 2019), and neuropsychiatric disorders (Sherwin, Sandhu, Dinan, & Cryan, 2016; Tetz & Tetz, 2016). Despite the large proportion and diversity of phages in the gut microbiome, the impact of phages on the bacterial community remains poorly understood as bacterial species have been the main focus of microbiome studies.

Different novel phage types have been detected as new detection methods are introduced, highlighting bias in previous research methods (Al-Shayeb et al., 2020; Serwer, Hayes, Thomas, & Hardies, 2007). These include for example large phages, aggregating phages, and phages that infect unculturable bacteria. Therefore, it is important to be able to study all types of phages in a community, in order to properly understand their role and impact on the microbial ecosystem. This too, requires novel methods.

Metagenomic analyses of complex microbial communities have emerged over the last two decades revealing a large proportion of microbes that were previously undetected, including bacteria, viruses, and archaea (Breitbart et al., 2008; Brown et al., 2015; Edgar et al., 2022; Lagier et al., 2012). Metagenomics are less biased against phages of different shapes and infection-patterns, as culturing is not required. The possibilities of comparative genomics and genome assemblies from metagenomic data have thus led to an increase in novel species identification of bacteria, archaea, and phages. A lot can be learned from the genome sequence of novel species, but phenotypic characterisation and gene functions are still largely culture-dependent. However, the road from *in silico* genome to *in vitro* isolation is long and complicated.

Successful isolation of microbial species discovered exclusively in metagenomes is rare, yet it has been achieved (Bomar, Maltz, Colston, & Graf, 2011; Browne et al., 2016; Pope et al., 2011).

Few phages have been isolated based on metagenomics sequencing data. Phages are dependent on the availability of a specific bacterial host for replication, as well as several other biological factors that are needed for their survival. Their genomes, however, are extremely diverse and leave little information about hosts, optimal growth conditions, and alternate life cycles. Over the past decade, a series of webserver tools for phage-host prediction have emerged based on sequence similarity, CRISPR spacer matches, and similarity to known phage-host pairs (Edwards, McNair, Faust, Raes, & Dutilh, 2016), including; PHERI, HostPhinder, Host Taxon Predicter, and WIsH (Baláž, Kajsík, Budiš, Szemeš, & Turňa; Gałan, Bąk, & Jakubowska, 2019; Galiez, Siebert, Enault, Vincent, & Söding, 2017; Villarroel et al., 2016). These tools can provide valuable methods for narrowing down potential host species, but their efficiency is limited to well-studied phage-host pairs and well-characterised host species. As a majority of microbial species are yet unknown (Staley & Konopka, 1985), and phage culturing is also subject to environmental conditions (Serwer, Hayes, Thomas, Demeler, & Hardies, 2009; Serwer et al., 2007), *in vitro* isolation of phages that are discovered in metagenomics presents a great challenge.

One example of successful *in vitro* isolation of a metagenomics-derived phage is the CrAss-phage (Dutilh et al., 2014). When the CrAss phage genome was discovered in 2014 in human gut microbiome sequences, it was shown to occupy up to 90% of the human gut viral community (Dutilh et al., 2014). Despite the abundance of this novel phage, it was not until 2018 a member of the CrAss-phage family was first isolated (Shkoporov et al., 2018) and to this date only four members of the CrAss phage family have been isolated (Guerin et al., 2021; Hryckowian et al., 2020; Shkoporov et al., 2018). Both φCrAss001 and φCrAss002 were shown to behave differently than expected in culture, producing no visible plaques on the propagating host (Guerin et al., 2021; Shkoporov et al., 2018). This altered life cycle have likely contributed to the difficulty in isolation of these and other phages using traditional methods and should be considered when trying to isolate *in silico* phages.

As more phages are discovered *in silico*, it has become more apparent that traditional methods of phage isolation are biased towards smaller phages with known life cycle traits (Serwer et al., 2007). However, phages with large genomes of >200.000bp (jumbophages) are also found in metagenome studies despite the underrepresentation of sequenced and isolated large phages (Devoto et al., 2019; Yuan & Gao, 2017) (Al-Shayeb et al., 2020). Phage genomes are packed tightly to maximise genetic storage in the small capsids. Phages with larger genomes are thus likely to have a larger capsid, which might alter the behaviour of some of these phages *in vitro*. Traditional approaches, such as the plaque assay, will not be suitable for the larger phage isolations without modification, as diffusion of the large particles is inhibited by the dense agar matrix (Serwer et al., 2007). As *in silico* organisms have proven difficult to isolate, the challenge lies in studying and characterising important phages in a culture-independent method. Several approaches have been suggested including microfluidic PCR (Tadmor, Ottesen, Leadbetter, & Phillips, 2011), viral tagging (Deng et al., 2012) and phageFISH (Allers et al., 2013).

All three methods can aid in phage host identification independently of culture, and can potentially answer questions about the host range of uncultured phages. However, microfluidic PCR and viral tagging do not allow characterisation of viral behaviour or visual detection.

PhageFISH can potentially characterise phages without the need for culturing (Allers et al., 2013). PhageFISH utilises the known phage genome to produce several long DNA probes that are specific to each phage for fluorescent detection and observation along with bacterial probes. The method was developed for improved characterisation of phage-host interactions in marine systems. Allers et al. were able to detect intra- and extracellular phage DNA, as well as perform host identification and quantification. It was also possible to study different waves of infection and discriminated between early and late infection stage (Allers et al., 2013). As the method uses specific phage probes designed from the phage genome, phageFISH could potentially be employed for uncultured *in silico* phage detections in metagenomics samples, including faecal microbiomes.

In this study, we employed the phageFISH method (Allers et al., 2013) to visualise five jumbophages *in silico* that were previously assembled from Danish pig faecal metagenomes (used in Al-Shayeb et al., 2020) thereby confirming that they are true phages. PhageFISH allows visualisation directly in the native faecal matrix via specific DNA probes. To our knowledge, this is the first time phageFISH has been used to detect and visualise previously uncultured jumbophages directly in faecal samples, linking *in silico* phage genomes to *in vitro* visualisation.

## Material and Methods

### Sample collection

Pig faecal samples were collected as part of VetForligII project from farms around Denmark from 2014 to 2016. Faeces were stored at -80°C until use.

Five novel large phage genomes were assembled based on sequence data from the metagenomes 10 pig faecal samples. Each phage genome was based on similar sequence assemblies from two different faecal samples. The novel phages are denoted Phage A, B, C, D, or E (Supplementary Table S1).

### Isolation of faecal large viral population

Large viral population was obtained from faecal sample as described previously by Saad et al. 2019 with modifications. Briefly, 1 gram frozen faecal matter was suspended thoroughly in 40ml SM-buffer (0.1M NaCl, 10mM MgSO_4_, 50mM Tris-HCl pH 7.5, 0.01% (w/v) gelatin). The suspension was shaken at 800rpm overnight at 4°C to dissociate phages. The suspension was then centrifuged at 5000x*g* for 10 minutes to pellet large bacteria and debris. The supernatant was then centrifuged at 15.000x*g* for 1 hour at 4°C to pellet large phages. The pelleted phages were resuspended in 5ml SM-buffer, and the centrifugation process was repeated before adding an equal volume of chloroform to remove bacteria. The final viral population was centrifuged at 15.000x*g* again and suspended in 2ml SM-buffer.

### Large viable phage detection with plaque assay

Standard plaque assays were adapted for large phage isolation as described by Saad et al. 2019 with few modifications. Two strains of *Escherichia coli*: *E*.*coli* 11303 and *E. coli* BAA-1025, were grown on lysogeny broth (LB) agar plates overnight and 3-4 colonies were inoculated into LB broth and incubated for 2-3 hours to reach exponential growth phase. 250μl of the exponential phase bacteria and 100μl viral population were co-incubated for 10 minutes before adding 5ml LB broth (or BHI broth) with 0.35% agar supplemented with 10mM CaCl_2_ and 10mM MgSO_4_. The bacteria-phage mix was immediately poured on LB (or BHI) agar plates. All plates were inoculated in 4 replicates and incubated at 30°C or 37°C as well as aerobically or anaerobically, for 24-48 hours.

### DNA extraction

DNA was extracted from each faecal large viral population using Blood and Tissue kit (QIAGEN, Denmark). The large viral population sample, isolated as described above (Isolation of faecal large viral population), was centrifuged at 15.000x*g* for 1 hour. Pellet was resuspended in 200μl ATL buffer and 20μl proteinase K, and lysed overnight at 56°C with shaking. Protocol “Purification of Total DNA from Animal Tissues” was followed afterwards according to the manufacturer’s instructions. Samples eluted in 2×50μl preheated AE buffer and DNA concentrations were measured with Qubit (Invitrogen).

### Primer design and probe synthesis

Several DNA primers amplifying a 300bp fragment (Supplementary Table S2) of the targeted phage genomes were designed for each phage (Supplementary Table S3). DNA probes were synthesised using the designed primers followed by PCR DIG-probe Synthesis kit (Roche) according to the manufacturer’s instructions in 50μl reaction volumes. A touchdown reaction cycle was used for PCR product amplification (94°C 4min, 35 cycles of {94°C 30s, 60°C 30s (−0.20°C/cycle), 72°C 30s}, 72°C 4min).

45μl PCR product was purified with Gene Clean Turbo for PCR kit (MP Biomedicals) according to the manufacturer’s instructions. Probes were eluted in 30μl sterile nuclease-free water. The PCR product size, concentration, and integrity were evaluated on Bioanalyzer 2100 (Agilent).

A single unlabelled probe from each sample was sequenced with Sanger sequencing to confirm the probe sequence (Eurofins).

### Detection of phages across faecal samples

The specific phages were detected across all faecal samples included in this study (Table 1) by PCR amplification using the same primers that were also designed for probe synthesis with the same conditions described above.

**Table 1:** Detection of assembled phage genomes in extracted viral population from each faecal sample by PCR amplification.

| DNA | PRIMER | A5 | A6 | B3 | B4 | B7 | C5 | D3 | D6 | E1 | E2 | E3 | E4 | E5 | E6 | E7 |
| --- | --- | --- | --- | --- | --- | --- | --- | --- | --- | --- | --- | --- | --- | --- | --- | --- |
| **A** | F12 | + | + | - | - | - | (+) | - | - | - | - | - | - | - | - | - |
|  | F33 | + | + | (+) | - | - | + | + | - | - | - | - | - | - | - | - |
| **B** | F42 | + | + | + | + | + | + | + | - | - | - | - | - | - | - | - |
|  | F71 | + | + | + | + | + | + | + | + | - | - | - | - | - | - | - |
| **C** | F67 | - | - | (+) | - | - | + | - | - | - | - | - | - | - | - | - |
|  | F78 | + | + | - | - | - | + | - | - | - | - | - | - | - | - | - |
| **D** | F95 | + | + | (+) | - | - | - | + | + | - | - | - | - | - | - | - |
|  | F101 | + | + | + | + | + | - | + | + | + | + | + | + | + | + | - |
| **E** | F49 | + | + | - | - | - | - | - | + | + | + | + | + | + | + | - |
|  | F59 | - | - | - | - | - | - | - | ND | + | + | + | + | + | + | + |

<sup>+</sup>Single band at 300bp
<sup>(+)</sup>Faint band at 300bp
- No band, smudged bands, or inconclusive data
<sup>ND</sup>No data

### PhageFISH

Fluorescence *in situ* hybridisation of phages was adapted from Barrero-Canosa and Moraru 2019 (Barrero-Canosa & Moraru, 2019). All buffers are listed in Supplementary Table S4.

A 10μl loop-full of frozen faecal matter was suspended in 50μl PBS. 10μl of the faecal suspension were placed directly on a poly-L-lysine coated glass slide and smeared into a thin layer. The smeared sample was dried for 30 minutes at room temperature before fixing in 500μl of 1% paraformaldehyde (v/v) for 1 hour at room temperature. Slides were rinsed in PBS to remove paraformaldehyde solution and moved to a permeabilisation buffer for 1 hour on ice. The slides were gently rinsed in PBS and sterile water before incubation in 10mM HCl for 10 minutes. Slides were rinsed again in PBS and sterile water, and dried in 96% ethanol.

16S rRNA probes for bacteria were hybridised by incubation in hybridisation buffer I for 3 hours at 46°C. After hybridisation, slides were incubated in wash buffer I at 48°C.

Phage probes were hybridised to glass slides according to Barrero-Canosa & Moraru 2019 (SI methods).

For mounting and DAPI DNA-universal staining, slides were thawed for 10 minutes at room temperature and 10μl SlowFade Gold with 5μg/ml DAPI were added to the slides and covered by 24×50mm cover glass. Samples were sealed with clear nail polish and visualised with fluorescence microscope (Olympus BX53, PlanApo N 60x objective).

Sample phageFISH images were captured using CellSense software (Olympus) and processed in ImageJ.

## Results

### Isolation of large viral population

Large phages were isolated from each faecal sample for DNA purification and probe synthesis. The methods were evaluated based on phage recovery by plating phages on *E. coli 11303* and *E. coli BAA-1025* in dilute LB agar. Temperature variation had no observed effect on plaque formation in our assays. All tested large viral populations were able to produce plaques on the *E. coli* strains. Variation in morphology of the dominating plaque types could be observed on the plates suggesting presence of several different phage types (Supplementary Figure S1) (Saad et al., 2018).

### Detection of phage presence across faecal samples

To confirm the presence of the assembled phage genomes from our metagenomics samples (targeted phage A-E), a PCR reaction was performed to detect the unique probe fragments in the samples (Table 1). This further confirmed the successful extraction of large phages in the population samples and indicated Phage A-E presence in each sample using PCR only (Table 1). Interestingly, Phage A appears to be present in almost all the tested samples except F67 (initial origin of Phage C – Table 1) and F59 (initial origin of Phage E – Table 1), whereas Phage E was found less frequently.

### Probe synthesis

A DNA target region was selected for each phage for phageFISH detection, and 34 primer pairs were designed to amplify the targeted regions (Supplementary Table S3), 15 pairs were successful at amplifying the targets (Supplementary Table S3). Probes were synthesised from the amplified regions where the primer pairs were successful and for each of the five phages. Seven probes belong to Phage E. For Phage C only a single probe could be synthesised, Phage A and D two probes were synthesised, and for Phage B three probes were obtained (Table 1).

### PhageFISH set-up

Auto-fluorescence in the 475-650nm emission spectrum (FITC filter) is common in faecal samples (Dinoto et al., 2006) and was evaluated by comparing one faecal sample (F12) stained with DAPI only (Supplementary Figure S2a) to samples probed with either target or non-target probes (Supplementary Figure S2b and S2c). Probe group E was used for the comparison on sample F12 (no Phage E targets, Supplementary Figure S2b and Table 1) and sample F49 (Phage E target) (Supplementary Figure S2c, and Table 1). Auto-fluorescent debris is detected in the FITC filter spectrum image of the non-probed faecal sample as dim green signals (Supplementary Figure S2a and S2b). Phage signals, on the other hand, are more distinguishable and defined as individual intense dots (as also described by Allers et al.) or as intense signals in the shape of bacterial cells (Supplementary Figure S2c).

Bacterial cells were probed with cyanine dyes (Cy3 and Cy5) integrated into the probes before hybridisation. Using cyanine-labelled probes instead of HRP-labelled probes amplified by CARD shortened the protocol and allowed for visible signals from the bacterial probes. Signals produced by cyanine-labelled probes are not affected by the phage probe hybridisation procedure and the nonEUB338-Cy5 probe could be used as a negative control (Supplementary Text).

All probe groups were able to produce a phage signal in at least one sample (Table 2). To assess the impact of using multiple probes in a sample, the probe groups were compared (Supplementary Figures S3a-e), but no obvious difference in signal intensity or abundance was observed between samples that are probed with 1 or 7 probes. This allowed for visual detection of never before isolated phages with unknown hosts directly in the faecal material. Phage detection quality appears highly dependent on the faecal sample and the cell distribution in the faecal smears as witnessed by visual inspection of approximately 140 smears.

**Table 2:** Co-localisation detected in individual faecal samples by phageFISH in faecal smears, free phages and advanced infection could be detected in all samples.

| Phage group | Faecal sample | Free phage | Early infection | Advanced infection | Burst |
| --- | --- | --- | --- | --- | --- |
| A | F12 | Yes | Yes | Yes | Yes |
| A | F33 | Yes | Yes | Yes |  |
| B | F42 | Yes |  | Yes |  |
| B | F71 | Yes |  | Yes |  |
| C | F67 | Yes | Yes | Yes | Yes |
| C | F78 | Yes | Yes | Yes |  |
| D | F95 | Yes | Yes | Yes | Yes |
| D | F101 | Yes |  | Yes | Yes |
| E | F49 | Yes | Yes | Yes | Yes |
| E | F59 | Yes |  | Yes | Yes |

### PhageFISH

Bacterial probe EUB338-Cy3 was hybridised to bacterial cells to study the co-localisation between phage signals and bacterial signals. Co-localising signals indicate phage adsorption and infection of the targeted organisms, and further support successful hybridisation to phages. For Phage C and D, phage signals do not appear to co-localise well with the bacterial signal (Figure 1a and Supplementary Figure S4e-h). Using DAPI dye to visualise all DNA (including fungi, protozoa, mammalian cells) showed better co-localisation (Figure 1b). Figure 1b also shows a tendency for phage particles to co-localise and have high affinity to the sample debris.

**Figure 1.**
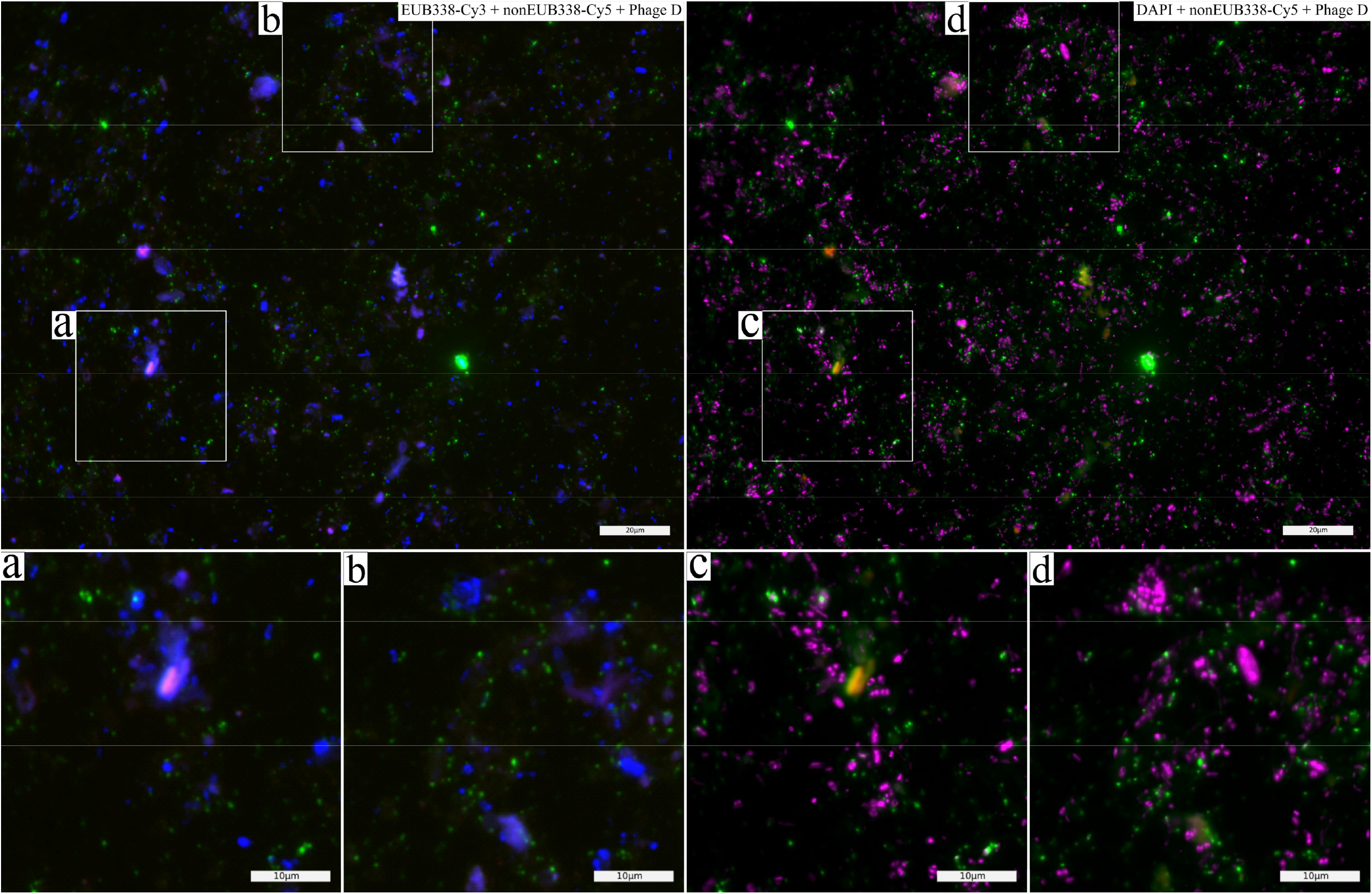
Co-localisation of phage signals to EUB338-Cy3 and DAPI signals. Microscopic image of faecal smear (sample F95) hybridised with bacterial probe EUB338-Cy3 (blue, only shown in a and b), and phage group D probes (signal simplified - green), and non-specific probe nonEUB338-Cy5 (negative control - red). All nucleic acids stained with DAPI (magenta, only shown in c and d). Co-localisation with phage signals is better observed with DAPI staining than with the bacterial probe EUB338-Cy3. Scale bars indicate 20μm and 10μm respectively..

It was possible to observe free phages and examples of advanced infection in all samples. Examples of early infection could also be observed in samples F12, F33. F78, F95, and F49, and examples of burst could be observed in samples F12, F67, F95, F101, F49, and F59 (Table 2). This is illustrated by sample F49 stained with Phage E probes (Figure 2a and 2b). The blue bacterial 16S rRNA probe signal co-localises with the green phage signal and represents several steps in the phage life cycle in one image; free floating, initial infection, propagation, and burst (Figures 2a and 2b). Examples of co-localisation observations are marked in the images of all samples available in supplementary material (Supplementary Figure S4).

**Figure 2.**
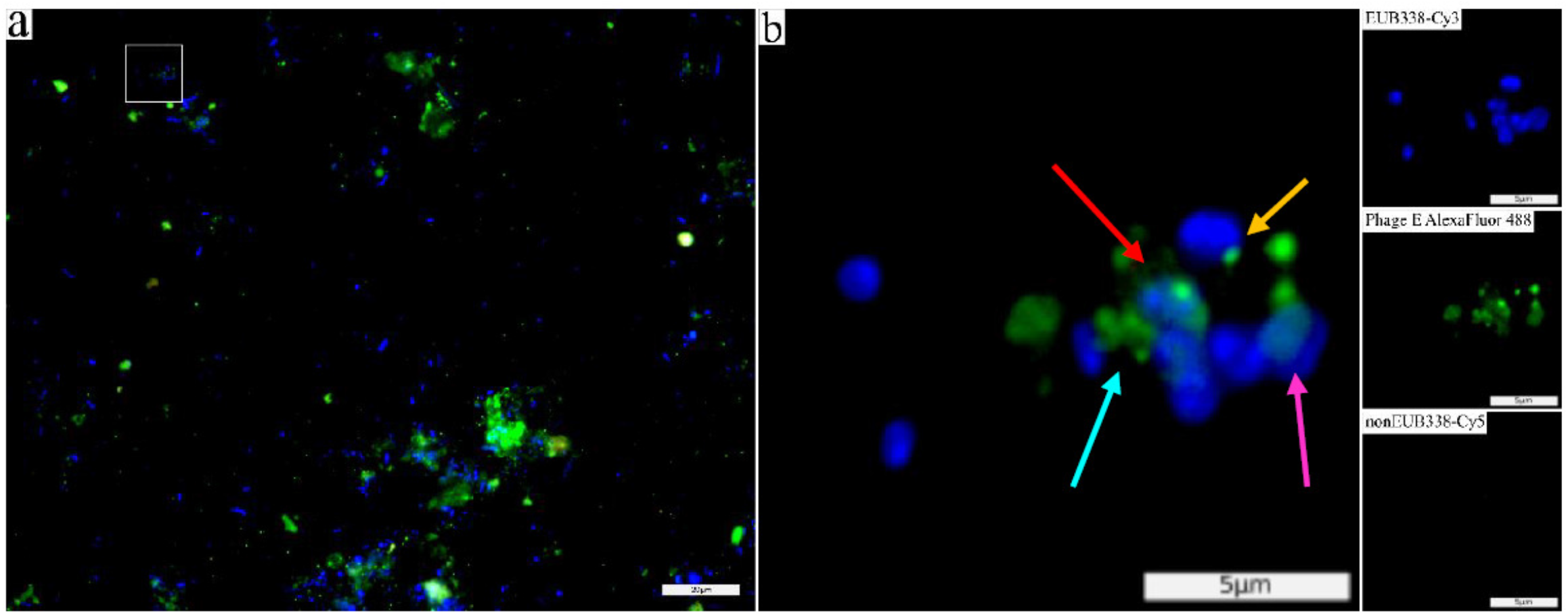
Co-localisation of phage signals and EUB338-Cy3 signals allows visualisation of phage-host interaction and phage behaviour. Microscopic image of faecal smear (sample F49) hybridised with phage group E probes (green), universal bacterial probe EUB338-Cy3 (blue), and non-specific probe nonEUB338-Cy5 (red). Phage behaviour was observed as free phages (blue arrow), early infection (yellow arrow), advanced infection (red arrow), and burst (pink arrow). a) White box indicate enlarged area in b. Panels to the right show separated signals (EUB338-Cy3, Phage probe E, and nonEUB338-Cy5) of the enlarged area. Scale bars indicate 20μm (a) and 5μm (b).

## Discussion

Studying novel phages and phage types discovered with the rapidly increasing metagenomic research is paramount to our understanding of the ecosystems we investigate. Traditional methods of phage isolation have proven insufficient for isolation of many types of phages (Dutilh et al., 2014; Serwer et al., 2009) and new methods reducing this bias are necessary. Here, we attempted to confirm that five jumbophage genomes, only known *in silico*, were true phages using phageFISH directly in the natural environment of the community (faeces). This allowed visualisation of phage-host interaction in a culture-independent fashion for all five phages.

To reduce the bias against large phages, large viral particles were isolated from each faecal sample and DNA was extracted from the large viral population. DNA content isolated directly from faecal samples would likely be dominated by bacterial DNA (Stephen & Cumming. J. H., 1980), which further complicates the detection of phages using a synthetic probes approach. Traditionally, bacterial contaminants are removed by filtration. However, filtration also eliminates large phage particles, therefore a different method had to be adopted (Serwer et al., 2009). The method presented by Saad et al. was employed with minor modifications for phage extraction from faeces (Saad et al., 2018). As faeces are more microbiologically concentrated than sewage, it was important to optimise sample suspension and phage dissociation from debris to maximise the yield from the small faecal samples. It was possible to extract large phages from 0.5g of faeces.

In a previous study where geneFISH, the basis of phageFISH, was used, an approx. 40% detection efficiency for single copy genes using a single probe was estimated (Moraru, Lam, Fuchs, Kuypers, & Amann, 2010). Based on this, Allers et al. argue that single copy organisms (such as phages) are unlikely to produce signals strong enough for detection of single free phages and show that detection efficiency increased with each added probe using long probes and CARD signal amplification (Allers et al., 2013).

Here however, we found it possible to detect free phages (Phage C) using a single 300bp probe (probe C5 in Table 1, Supplementary Figure S3c). This could be due to the variation in the targeted phages between this study and that of Allers et al. The large genome of unknown phages here may behave differently than the small, well-studied phages by Allers et al. (Allers et al., 2013). A tendency for phages to adhere to debris in the sample may also affect individual probe detection, as signal strength may increase by accumulation (Supplementary Text).

As opposed to the protocol presented by Allers et al. and Barrero-Canosa & Moraru, cyanine-labelled probes were used for bacterial gene detection instead of HRP-labelled and CARD-amplified probes. Most bacterial species encode multiple copies of the 16S rRNA gene, making the cyanine fluorescent signal strong enough for detection in normal fluorescence microscopy. This was verified by the optimisation of each Cy-labelled probe in cultures before hybridisation in faeces. We tested CARD-amplification of bacterial genes and found great resolution of cells in culture, but the signals were quenched after phage-probe hybridisation (Supplementary Text).

Cy-labelled probes are a faster and cheaper alternative to DIG-labelled or HRP-labelled probes and met the standards necessary in this study. It also allowed us to use the negative control probe nonEUB338-Cy5 at the same time as both bacterial and phage probes. Furthermore, the nonEUB338-Cy5 probe acted as a control for background fluorescence and auto-fluorescence in the faecal smears. This was particularly useful when images were digitally enhanced for better recognition of phage signals and for distinguishing different stages of phage infection (Supplementary Text).

PhageFISH has previously been used to study phage-host infection dynamics (Dang, Howard-Varona, Schwenck, & Sullivan, 2015; Jahn et al., 2021). Here, we used a universal bacterial probe to visualise bacterial cells interacting with the phages. Co-localisation of bacterial and phage signals were observed in all samples, but were not common (Table 2, Figure 1 and Supplementary Figure S4). This could be explained by the targeted phages and their hosts not making up a large proportion of the sample as many bacteria and their phages are abundant in faecal samples (Stephen & Cumming. J. H., 1980).

The majority of phage signals did not co-localise with any bacterial signal, indicating a large number of free phages in the sample. As conditions change along the GI tract, phage adsorption may be inhibited or reduced, causing a large proportion free phages (Crespo-Piazuelo et al., 2018; Looft et al., 2014; Zhao et al., 2019).

Here, free phages were defined as a phage signal with no other signal overlapping. This definition depends on the efficiency of the DAPI dye and the eubacterial probe (EUB338, Supplementary Table S2), as phages infecting an unstained bacterial cell will result in an interpretation as free phage. DAPI is widely used for estimation of total cell numbers in faeces and is generally trusted to dye all nucleic acids in a sample (Dinoto et al., 2006; Harmsen, Raangs, He, Degener, & Welling, 2002; Selim et al., 2005). As DAPI-signals were often seen to co-localise to phage-signals better than the bacterial probe signal, there could be an issue with EUB338-Cy3 probe or the phages could infect a host untargeted by the probe.

The eubacterial probe EUB338 probe does not target archaea, fungi, and mammalian cells, all stained by DAPI. Furthermore, EUB338 has reduced affinity to certain bacterial species (Daims, Brühl, Amann, Schleifer, & Wagner, 1999), the majority of which are not typically associated with animal faeces. The untargeted phylum *Verrucomicrobia*, however, may be present in gut microbiomes and represented by the *Akkermansia* genus (Crespo-Piazuelo et al., 2019). As the relative abundance of *Verrucomicrobia* members in pig faeces is relatively low (Crespo-Piazuelo et al., 2018; Niu et al., 2015), it is unlikely that the detection of free phages with a single probe can be explained as infection of an unstained bacterial cell. This is further supported by the observations of advanced infection in bacterial cells in multiple samples (Supplementary Figure S4).

The EUB338-Cy3 probe may perform less efficiently in faecal smear than in the culture in which it was optimised. As bacterial cells in the faecal smear cannot be quantified independently of the bacterial probe, it was difficult to optimise the probe affinity in the faecal environment and unsuccessful binding of the bacterial probe could happen due to the physiological conditions of the faecal material. This may result in reduced co-localisation of signals. Furthermore, if conditions in the faecal sample at the time of the sampling are not optimal for infection, and the phages exist primarily as free phages, the chance of random co-localisation without infection would be higher for the DAPI stained than the EUB338-Cy3 labelled cells, simply due to the higher number of DAPI stained cells.

In the cases of co-localisation with the bacterial probe, different stages of the phage life cycle can be observed. This is particularly evident in one of the studied faecal samples (Figure 2). To our knowledge, this is the first example of phage-host interaction observed for a non-isolated phage in faeces with no known host.

As the numbers of *in silico* identified phage genomes become more abundant, (e.g. CrAss phages) it is necessary to find new methods of investigation. Isolation of organisms from *in silico* to *in vitro* is not a straightforward process, as a large percentage of sequenced bacteria cannot be cultured *in vitro* increasing the risk of the phage of interest being unculturable too.

The CrAss phage remains the best example of successful *in vitro* isolation of an *in silico phage* (Dutilh et al., 2014; Shkoporov et al., 2018). Only four members of the highly abundant phage have been isolated *in vitro*, φCrAss001 (Shkoporov et al., 2018), φCrAss002 (Guerin et al., 2021), and two discovered by coincidence (Hryckowian et al., 2020). For both cases of targeted isolation of crAss-like phages, only a single phage was isolated *in vitro* despite broad experimental set-ups. The crAss-phage family is heavily abundant in human gut microbiomes, and may easily infect one of the several unculturable species inhabiting the human gut microbiome (Le Chatelier et al., 2013) explaining the difficulty in isolation of these species. Furthermore, both φCrAss001 and φCrAss002 were shown to behave differently than expected in culture, producing no visible plaques on the propagating host (Guerin et al., 2021; Shkoporov et al., 2018), which could also affect the success of isolation using traditional methods. This could also be the case in our study and the general study of large genome phages.

In the 2019 study by Devoto et al., evidence was found of the unusually large Lak phage inhabiting the gut microbiome of several animals and humans with a genome size of >500kb. According to CRISPR spacer sequences, the phage infects a species of *Prevotella* but could not be isolated in culture (Devoto et al., 2019). *Prevotella* can be cultured *in vitro*, yet it is difficult to grow in confluent lawns of low-percent agar as is necessary for traditional phage isolation. This is evidenced by the low number of *Prevotella*-phage isolated in culture as well as our own experience in the lab.

The prospect of isolating the phage is complicated by the size of the genome. Relatively few phage genomes of similar size have been described and little is known of the ways in which the increased genomic real estate may influence the function of the phage and the isolation parameters necessary to accommodate the large phages (Hendrix, 2009; Yuan & Gao, 2017). Lak phage was attempted isolated with no special consideration of phage size, leaving little chance of isolating it in culture (Devoto et al., 2019).

Similarly here, all five phage genomes are based solely on metagenomic assemblies from pig gut microbiomes and are larger than the average phage (Hendrix, 2009). PhageFISH and other fluorescent techniques offer a method of observation in a culture independent fashion. Without any information about potential hosts, the phages were observed directly in their natural environment to achieve unbiased visualisation of the unknown phages.

In conclusion, the adapted phageFISH method presented here allows detection of *in silico* jumbophages in their natural environment regardless of their size, life cycle, and host organism. Further studies combined with family or species specific bacterial probes, may allow potential phage hosts identification irrespective of the bacterial host ability to be cultivated. As *in silico* phages have proven difficult to isolate *in vitro*, development of such methods is important to facilitate biological validation of novel phages, as well as characterisation in a culture-independent manor.

## Acknowledgements

The authors would like to thank Terje Svingen, DTU FOOD, for the help with the fluorescence microscopy. The authors would also like to thank the two reviews for their valuable comments and suggestions. This work was funded by the Novo Nordisk Foundation Grant NNF16OC0021856.

